# Robust Consensus Nuclear and Cell Segmentation

**DOI:** 10.1101/2025.04.03.646876

**Authors:** Melis O. Irfan, Eduardo A. González-Solares, Tristan Whitmarsh, Alireza Molaeinezhad, Mohammad Al Sa’d, Claire M. Mulvey, Marta Páez Ribes, Dario Bressan, Nicholas A. Walton

## Abstract

Cell segmentation is a crucial step in numerous biomedical imaging endeavors; so much so that the community is flooded with publicly available, state-of-the-art segmentation techniques ready for out of the box use. Assessing the virtues and limitations of each method on a tissue sample set and then selecting the optimum method for each research objective and input image is a time consuming and exacting task that often monopolizes the resources of biologists, biochemists, immunologists and pathologists; despite not being their project primary research goal. In this work, we present a segmentation software wrapper, coined `CellSampler`, which runs a selection of established segmentation methods and then combines their individual segmentation masks into a single optimized mask. This, so called ‘uber mask’, selects the best of the established masks across local neighborhoods within the image, where the neighborhood size and the statistical measure which determines the qualitative term ‘best’ are both chosen by the user.

## 1 INTRODUCTION

Segmentation is a prevalent challenge within medical imaging across a wide range of different modalities. Cell segmentation, in particular, enables the counting, characterization, and spatial mapping of distinct cell types. A single analysis can require cell segmentation of tens to hundreds of tissue samples; common practice being to select a single segmentation method that is seen to perform well on a few samples and then to assume that it will perform optimally on the full cohort. However, in reality, different segmentation methods may work better for different tissue samples. Within one sample alone, the best method may depend on the tissue region or cell type. Furthermore, even if time and staff resources would allow the option of manually annotating every tissue sample within a trial, Wiggins et al. (2024) have shown that variability between different expert annotators, or just the same annotator at different times, reduces the reproducibility of the subsequent analysis and scientific conclusions. In this work we aim to capitalizes on the strengths of multiple segmentation methods through the use of a consensus technique.

Medical image segmentation methods typically rely on an initial set of human annotated reference images: either as training samples for deep learning algorithms (Schmidt et al., 2018; Greenwald et al., 2022; Xiao et al., 2021; Stringer et al., 2021) or as Bayesian priors for maximum likelihood methods (Warfield et al., 2004). Artaechevarria et al. (2009) discuss the use of *a priori* human annotated segmentation masks, referred to as ‘atlas’ images, within the use of expectation–maximization algorithms and demonstrate that not only do multiple atlas images improve segmentation results for brain MRI data but so does altering the segmentation strategy across localized regions of the full image.

Recent developments in image processing have placed researchers with a requirement for nuclear segmentation in the fortuitous position of having numerous, proficient algorithms available to them. The challenge now is how to aptly select which technique to employ for each region of interest. We introduce a consensus segmentation technique capable of considering the nuclear segmentation of any number of individual segmentation algorithms. This consensus technique determines which algorithm outperforms all others within user-defined subsections of the image and combines the optimum segmentations from different techniques into a single choice result for the image, which we refer to as the uber mask.

Consensus segmentation is not an unusual approach within computer vision for optimizing boundary detections. However, the majority of applications only require a per-pixel approach to voting as they are intended for use on continuous image data. Boiangiu and Ioanitescu (2013), for example, detail a pixel co-association matrix approach which successfully combines segmentation algorithms to produce clearly segmented representations of people, animals, scenery etc. For per-pixel consensus voting, such as the one proposed in Boiangiu and Ioanitescu (2013), there is the inherent risk that several pixels within a single feature will be identified as different to their neighbors. In the simple example of an apple sitting on a table in front of a wall, the segmentation might clearly show the apple, the table and the wall as having clear boundaries but within any-one of those features there may also be additional, segmented regions due to the range of pixel intensities across a single feature. Whilst this is not an unacceptable feature of continuous image processing, it would prevent further analysis within cell segmentation. If two cells were erroneously identified as occupying the same spatial area by a nuclear segmentation algorithm then this would mislead any further interpretations of the tissue sample results.

For the specific case of single-cell nuclei detection, a consensus segmentation algorithm is required which maintains individual cell structures. Nguyen and Le (2022) move beyond the standard per-pixel analysis and implement a consensus vote which acknowledges the shape of the cell detected. In this work we extend their approach to be able to consider complex cell segmentation masks which include multiple features, as opposed to the binary masks required for the blood vessel segmentation show in Nguyen and Le (2022). In section 2 we present an overview of our segmentation combination pipeline and introduce the publicly available segmentation algorithms we intend to apply our consensus vote to. Within this section we also introduce various statistical criteria which can be used to construct the optimum consensus mask; a strength of our pipeline being that the user has the ability to select different criteria for different image sets based on their expertise and any cursory checks of the input images. Our pipeline is a general method applicable to any segmentation and in section 3 we present our results on a series of Image Mass Cytometry images. In section 4 we conclude.

## 2 MATERIALS AND METHODS

### 2.1 Overview of the Pipeline

Numerous segmentation algorithms are currently available for detecting cell boundaries within medical images; each with varying degrees of performance for different cell morphologies. The `python` pipeline presented in this work, `CellSampler`, is a comparison and combination wrapper which aims to capitalize on recent advancements within nuclear segmentation by employing a selection of state-of-the-art segmentation algorithms and combining their results into a single, optimized mask of cell nuclei.

In this work we explore the functionality of five established segmentation tools: Cellpose (Stringer et al., 2021), Mesmer (Greenwald et al., 2022), Stardist (Schmidt et al., 2018; Weigert et al., 2020; Weigert and Schmidt, 2022), a Watershed segmentation and an in-house implementation of DICE-XMBD (Xiao et al., 2021) followed by a Watershed segmentation. As Cellpose, Stardist and Mesmer have been optimized to perform on normalized input data, each input image is normalized between its 1^st^ (*d*_min_) and 95^th^ (*d*_max_) percentile values:

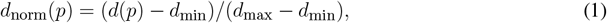

and then clipped so that any values less than zero/greater than one are set to zero/one. Lastly, local histogram equalization^1^ is applied to the scaled data to increase the image contrast.

### 2.2 Cellpose

Cellpose is a deep learning algorithm which employs convolutional neural networks (CNN) with the U-Net architecture to perform nuclear and cell segmentation. Cellpose has been trained on a variety of datasets and can also be retrained on specific images provided by the user. In this work we use `Cellpose3` and the built in ‘nuclei’ model type. Cellpose also allows the user to input an expected cell diameter and so we run two versions of Cellpose, one with an expected cell diameter of 6 pixels and one with an expected cell diameter of 10 pixels. In place of having to choose a single algorithm input parameter, such as cell diameter, `CellSampler` allows the user to try out several options simultaneously.

### 2.3 Mesmer

Mesmer is a segmentation algorithm which exploits the ResNet-50 CNN architecture and is trained on the TissueNet dataset (Barshir et al., 2012). In this work we use Mesmer for nuclear segmentation (as opposed to whole cell segmentation) and alter the image micron per pixel parameter to optimize our results. This scaling factor is a user input to the algorithm; Mesmer itself provides the scaling functionality. We use scaling factors of both 0.5 and 0.75*µm* throughout; resulting in two implementations of Mesmer.

### 2.4 Stardist

As with Cellpose, Stardist makes use of U-Net CNNs for nuclear and cell segmentation but, uniquely, enforces that the shape of each cell and nucleus can be characterized as a star-convex polygon.

### 2.5 Watershed

Our implementation of the Watershed detection algorithm uses a Voronoi-Otsu binarization to separate the nuclei from their background, finds the local maxima within the image as function of distance to the background and finally uses the watershed functionality from `scikit-image` to determine individual nuclei from these maxima markers. For optimum performance we first preprocess the image by removing any pixels with a magnitude 10 times larger than the median values of their 5 nearest neighbors. We also subtract off the background intensity from each circular region in the image of radius 75 pixels. The background intensity level is determined by smoothing the circular region pixels with a Gaussian beam.

### 2.6 DICE-Watershed

As a possible improvement to the standard watershed detection we also implement DICE-XMBD as a technique to segment the data before performing the watershed detection. This means that instead of performing watershed detection on the nuclear channel we instead do so on the DICE-XMBD probability maps which delineate each cell nucleus and cytoplasm as well as the slide background. DICE-XMBD requires both a nuclear and a cytoplasm channel to work from and, as some imaging data are not high enough resolution for or not specifically stained for cytoplasm detections, `CellSampler` provides a ‘synthetic’ cytoplasm channel.

The synthetic cytoplasm channel is formed by smoothing the nuclear channel image with a Gaussian and then using a Canny edge detector to find the edges meant to represent the cell detection. The nuclear and synthetic cytoplasm channel are combined and then smoothed again (to prevent any dips in intensity between the nucleus and cytoplasm edges) before being used as an input for DICE-XMBD. As we intended to implement the same U-NET architecture as DICE-XMBD, run on the same trained model as DICE-XMBD, we ensured that our combined nuclear and cytoplasm data were preprocessed (normalization and hot pixel removal) according to the DICE-XMBD methodology.

### 2.6 Consensus segmentation

Our consensus segmentation is an extension of the consensus approach proposed in Nguyen and Le (2022); tailored to fit the specific task of cell segmentation. The following is an overview of the Nguyen and Le (2022) algorithm:

1. The local neighborhood of a single pixel is defined as being within *s* pixels of the 2D pixel position (*x, y*) e.g. [*x − s* : *x* + *s*, *y − s* : *y* + *s*].
2. The Jaccard score is then calculated within the local neighborhood between all the contributing nuclear segmentation algorithms. The total Jaccard score for a single method is the sum of all the Jaccard scores between that method and the others.
3. The algorithm with the lowest total Jaccard score is eliminated from the process and then, in a second round of voting, the new total Jaccard score is recalculated for the remaining contributing algorithms.
4. A value is assigned to the pixel based on the algorithm with the highest total Jaccard score.
5. Steps 2, 3 and 4 are repeated for every pixel within the 2D image.

We choose the minimum over maximum interpretation of the Jaccard score (*J*) between two segmentation masks (*S*_*m*_ and *S*_*n*_) in our implementation:

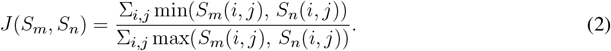

The kind of segmentation performed in Nguyen and Le (2022) is, however, always binary (segmentation masks of only zeros and ones) meaning that this is again a simplification of the problem the `CellSampler` consensus voting algorithm is trying to solve. For binary segmentation the task would be to determine tissue samples from a background of glass, in our case we need to be able to tell individual cells apart from each other as well as identifying them as being distinct from the background. Additionally, we want the metric for defining an optimum segmentation to be defined by the user based on a quick visual inspection of a few tissue samples. Chen and Murphy (2023) define numerous metrics with which to evaluate the success of a potential segmentation when no ground truth is know. Two of these metrics are 1) the number of cells detected and 2) the reciprocal of the natural logarithm of 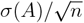 where *σ*(*A*) is the standard deviation of cell areas and *n* is the the number of cells. If the user had prior understanding of the number or consistency in size of the cells expected then these two metrics would be valuable additions to the Jaccard score. Therefore, we propose the following alterations to the Nguyen and Le (2022) algorithm:

1. The local neighborhood of a single pixel is defined as before, with *s* set to 40 pixels.
2. The ‘metric of merit’ is then calculated within the local neighborhood between all the contributing nuclear segmentation algorithms. The choices of metric are
  a. the total Jaccard Score (in order to compare the different nuclear masks all nuclear detection are converted to binary masks),
  b. the number of unique nuclei,
  c. 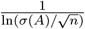
3. For the algorithm with the highest metric of merit we select those nuclear detections which have their (*x*,*y*) pixel center located within the neighborhood and save those nuclei to an array which stores the nuclear pixel locations alongside which method identified this nucleus.
4. Steps 2 and 3 are repeated for every neighborhood within the 2D image.
5. The array of all the saved nuclei from Step 3 is then checked for overlaps. Overlaps are first suggested as occurring when more than one nuclei centers are closer to each other than the radius (assuming nuclei areas are the area of a circle) of the largest nuclei in the overlap group. All suggested overlaps are then checked in the original segmentation masks to see if an overlap truly occurs. If more than one methods suggests a nucleus in the same location on the image (more than 3 pixels overlap) then this is classed as an overlap.
  a. In the case of an overlap, another neighborhood is defined around the overlapping nuclei. The metric of merit is again calculated between **all** the methods but this time the winner can only be picked from the selection of methods that contributed a nuclei to the overlap group in the first place.
6. After the removal of overlaps, the remaining nuclear detections form the uber mask.

The functionality used to determine nuclear detection pixel centers and areas is taken from the `skimage.measure` libraries.

### 2.8 Data

We make use of the publicly available BRCA2 dataset (Jackson et al., 2020) which provides Image Mass Cytometry (IMC) pathology images from breast cancer patients alongside ground truth segmentations created using CellProfiler (Stirling et al., 2021). Part of the `CellSampler` pipeline involves the use of a CNN trained on 2D data of size 512 by 512 pixels and so, to optimize the performance of this network, we require input images of size 512 by 512 pixels. As the BRCA2 dataset is significant in size (746 images), we can afford to select larger images and then just crop them to the 512 by 512 pixel area required for DICE-XMBD. We also expand our selection criteria to only include those images with over 400 cell detections as detected by the Watershed algorithm. This was done to ensure that the images processed contained a sizable number of cells to challenge the segmentation algorithms. The Watershed algorithm, as one of the most fundamental segmentation techniques, was used to provide the initial cell count check. These two requirements result in 258 images of 512 by 512 pixels (pixel size 1*µm*) with which to assess the nuclear segmentation algorithms.

We also show segmentations for two images provided by the IMC facility within the CRUK Cambridge Institute. Both images are from mouse tissue; one being lung tissue, the other being ovarian tissue.

## 3 EVALUATION

Within this analysis we use four scores to assess the success of each nuclear segmentation mask; the recall, precision, F1 and Jaccard scores. Each score ranges from 0 to 1, with 1 being the best possible result. These scores all rely upon a ground truth segmentation mask with which to assess possible detections as true positives (TP), fake positives (FP) and fake negatives (FN) against.

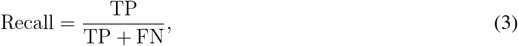

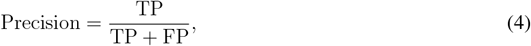

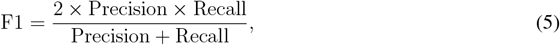

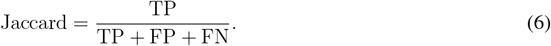

We rely on the Intersection over Union (IOU) measurement between a ground truth nucleus and predicted nucleus to decide how to classify each of the predicted nuclei as TP, FN or FP. For the example of a ground truth mask containing *N*_*o*_ nuclei and a mask predicted by a nuclear segmentation algorithm containing *N*_*p*_ nuclei, an IOU matrix of size *N*_*o*_ by *N*_*p*_ would be constructed. For computational ease, we do not calculate every element of the IOU matrix, instead choosing to assume that ground truth and predicted nuclei which have their centers (*x, y*) more than 80 pixels apart will not intersect and so will have an IOU value equal to 0.

First we assessed the predicted nuclei: all those with IOU values *<* 0.1 were designated FPs. A predicted nucleus with IOU values *>* 0.1, was marked as a TP match for the one ground truth nucleus with which it shared its highest IOU value. That ground truth nucleus would then be marked as matched. If the highest IOU (to two decimal places) for a predicted nucleus matched more than one ground truth nuclei then this predicted nucleus was marked as a merge error. Similarly if the highest IOU value for a predicted nucleus was matched to a ground truth nucleus that was already marked as matched then this would be a split error. Merge (one predicted nucleus matched to many ground truth nuclei) and split (multiple predicted nuclei matched to one ground truth nucleus) errors were both counted as FPs. Lastly we went through the ground truth nuclei and any of these which did not have an IOU value *>* 0.1 with a predicted nuclei were marked as FNs.

### 3.1 BRCA2

To represent the nuclear channel we use the average between two DNA marker channels: those stained with Iridium 193 and 191. Figure 1 shows ten examples of nuclear channel IMC images from the BRCA2 dataset; a variety of morphologies and intensities can be seen, even within this small subset. In Figure 2 we show the cell segmentations for a zoomed-in view of the first quarter of our first example image. As the ground truth segmentations provided are for the whole cell (not just the nucleus), we expanded our nuclear prediction masks each by a fixed number of pixels across all the images. We show the cell segmentation as red contours overlaid on top of the ground truth mask. Cellpose and Cellpose v2 refer to Cellpose used with a user specified cell radius of 10 pixels and 6 pixels, respectively. Mesmer and Mesmer v2 refer to Mesmer used with a user specified scaling factor of 0.75 and 0.5*µm*, respectively.

**Figure 1.**
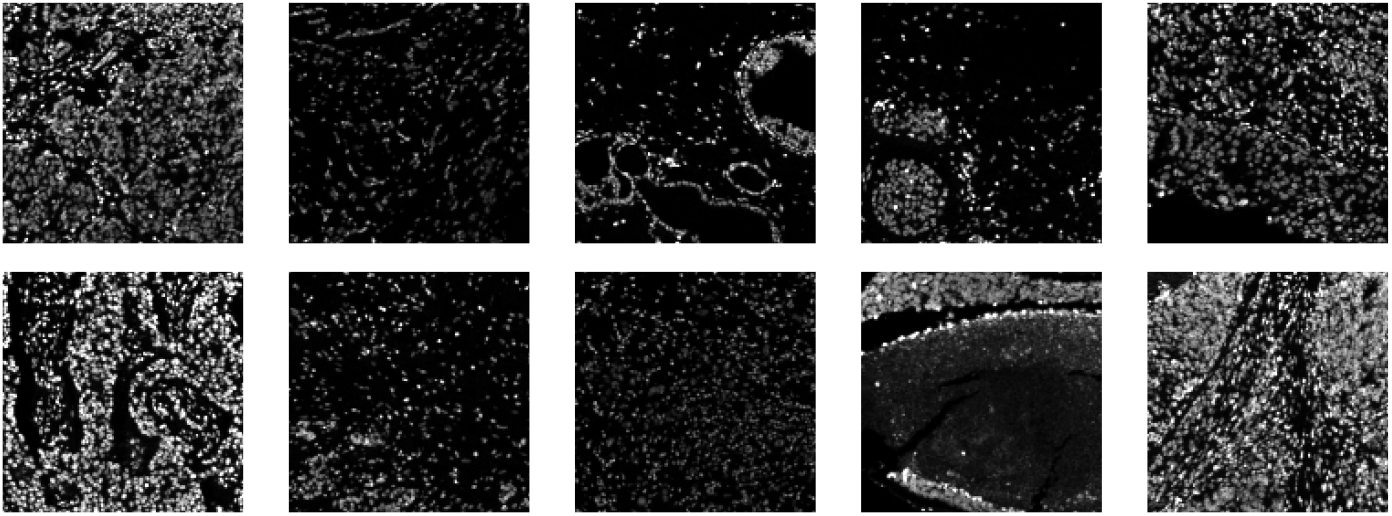
Ten example images (nuclear channel) from the BRCA2 dataset.

**Figure 2.**
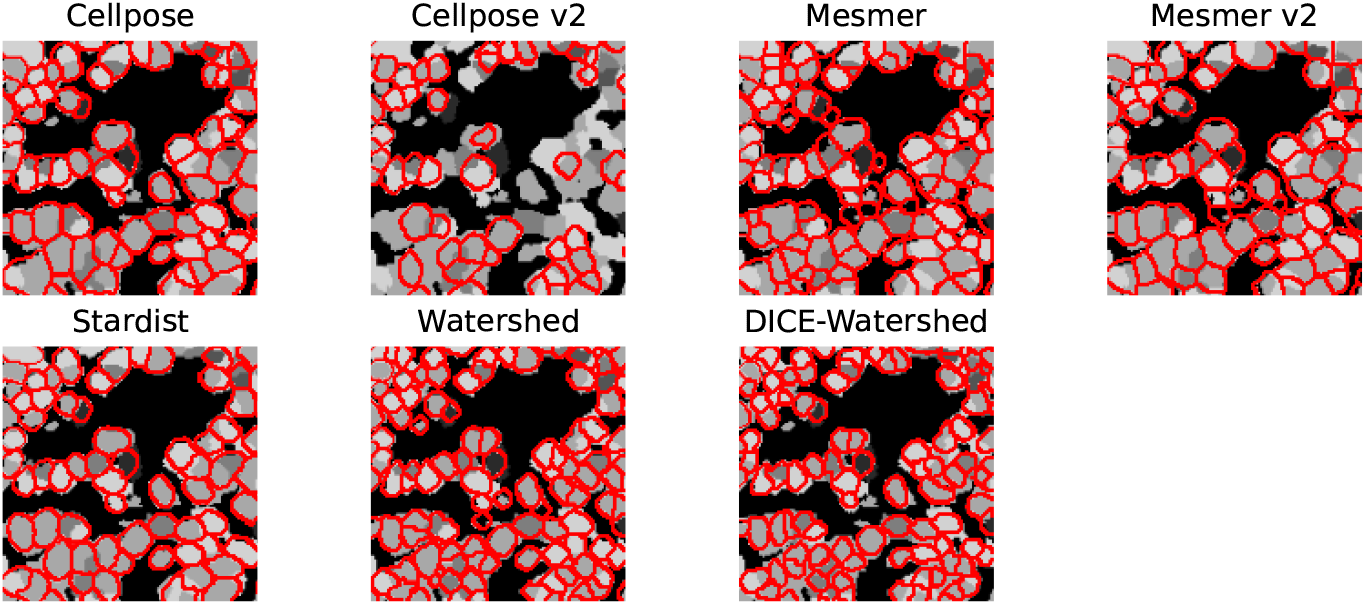
The segmentations produced by the five methods employed in this work overlaid onto the ground truth mask for a single example image. Only the first quarter (128 by 128 pixels) of the image is shown to highlight each distinct cell.

The top row of Figure 3 shows the cell detections within the uber mask for an example image. Each segmented cell is colored according the prediction algorithm that provided the segmentation. The three plots, from left to right, are the uber mask results for the three possible metrics of merit: 1) the reciprocal of the natural logarithm of the standard deviations of the nuclei areas, 2) the number of nuclei detected and 3) the Jaccard score. The bottom row shows the uber mask contours in red overlaid onto the ground truth mask. It is clear that the choice of metric makes a significant difference to the uber mask produced. While standard processes usually involves making an educated guess on which segmentation method to choose, the `CellSampler` pipeline allows the user to make an informed decision on what segmentation quality they would like to propagate into their data analysis from all the available methods. For the example shown in Figure 3, the user would be able to verify by eye that the uber mask produced using the criteria of number of nuclei detected best represented the ground truth. By examining the choice of masks for a couple of images and making a couple of ground truth annotations, it would be seen that under-segmentation is the key problem across the majority of regions and methods used in this analysis.

**Figure 3.**
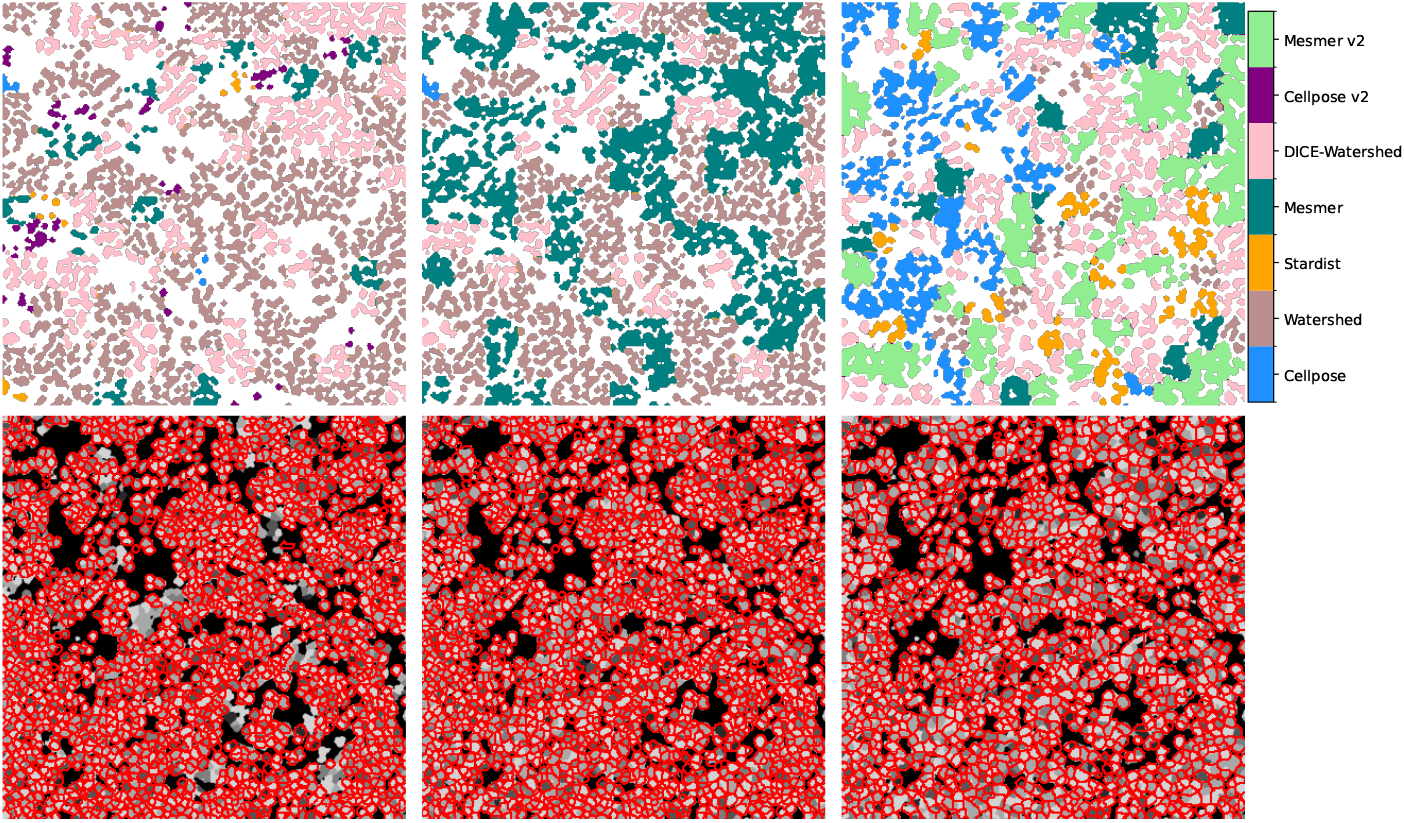
Top row: The cell detections selected from each method by the uber mask for our example image. Bottom row: the uber mask contours in red overlaid onto the ground truth mask. From left to right: the uber mask as made using 1) the reciprocal of the natural logarithm of the standard deviations of the nuclei areas, 2) the number of nuclei detected and 3) the Jaccard score as the score of merit.

For Figure 4 the recall, precision, Jaccard and F1 scores are calculated for all five segmentation algorithms as well as the three versions of the uber mask. The recall score is focused on how many true positives can be detected by an algorithm and here we see that the methods which make the largest number of cell detections overall perform the best. Cellpose v2 and Stardist, as used in their ‘vanilla’ mode i.e. without any specific training on this dataset, perform the worst having identified the fewest number of cells altogether. The problem of under-segmentation is perhaps one of the easiest to spot without the need for ground truth masks, through visual inspection of segmentation contours overlaid onto the nuclear image (such as those shown in Figure 2). The uber mask, version UM(2), which maximizes the number of nuclei detections has, unsurprisingly, the highest recall score. The precision score, however, is focused on how accurate an algorithm’s cell detections are and so will actively penalize methods which propose false positives. Conversely, we therefore see that the two methods which put forth the fewest cell detections have the highest precision as fewer total suggestions also means fewer false positive suggestions. The Jaccard and F1 scores are more comprehensive as they both include an assessment of true positives, false positives and false negatives. For both these scores we see that the Watershed, Mesmer and UM(2) uber masks have the highest scores, only differing from each other by 0.01 which is well within their error bars (shown on the lower plot of Figure 4) across the full dataset of 258 images. For all the methods we can see that, for the BRCA2 images, split errors (multiple predicted nuclei matching a single ground truth nucleus) are far more common than merge errors (a single predicted nuclei matching multiple ground truth nuclei).

**Figure 4.**
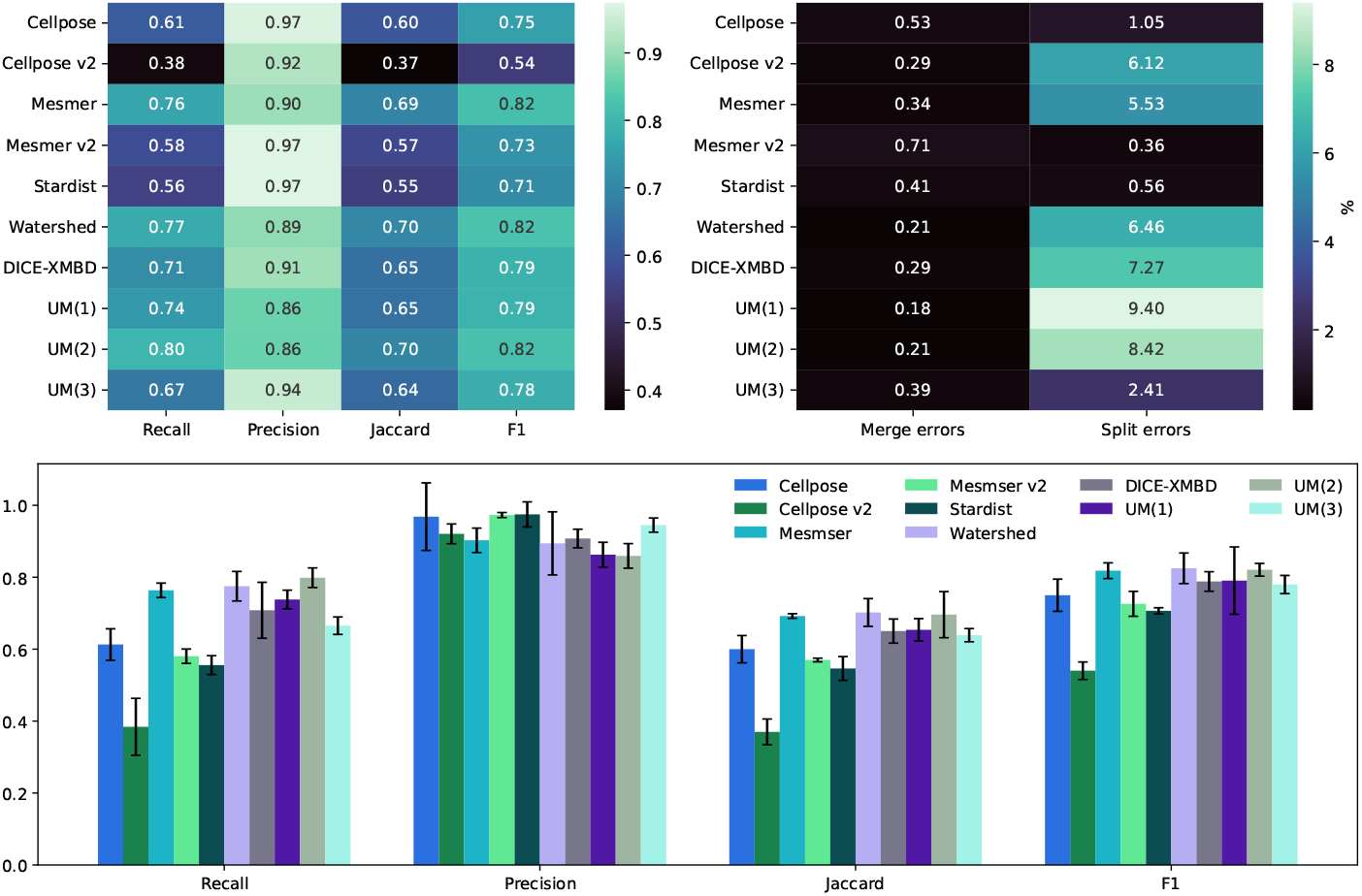
Top left: comparison between the median scores for all the nuclear masks created for the dataset of 258 images. Top right: the percentage of split and merge errors in our 258 images. Bottom row: the same as the top left but as a bar chart where the error bars are the mean absolute deviation values for the scores.

It is worth highlighting the fact that the metric of merit is not the only user parameter required for the creation of the uber mask; the user must also choose the size of the local neighborhoods within which to calculate the metric of merit. For the results shown in this section a pixel size of 40 was used to define the local neighborhoods. We also investigated the use of a neighborhood size of 20 and 80 pixels for the UM(2) uber mask; the scores for each pixel size are stated in Table 1. The scores are all consistent to within 0.01 no matter what neighborhood size is used, signifying that the uber mask is fairly robust to the choice of local neighborhood used as long as the choice is larger than the average expected cell radius. Splitting the image into neighborhoods smaller than a single cell would prevent the `CellSampler` pipeline from converging on a solution for a single given cell prediction.

**Table 1.** Recall, precision, F1 and Jaccard scores for the UM(2) uber mask using different pixels sizes to define the local neighborhoods.

| Pixel Size | Recall | Precision | F1 | Jaccard |
| --- | --- | --- | --- | --- |
| 20 | 0.79 | 0.86 | 0.69 | 0.81 |
| 40 | 0.80 | 0.86 | 0.70 | 0.82 |
| 80 | 0.79 | 0.87 | 0.70 | 0.82 |

### 3.2 Qualitative Results

In this section we take a more detailed look at the performance of the uber mask for two specific IMC images: one showing a section of the lung and one showing a section within ovaries. We have no ground truth annotations for these two IMC images, instead we assess the cell detections by eye using segmentations overlaid onto the input images. Figure 5 shows both the lung and ovarian IMC images. The lung sample was chosen for this analysis because it contained a metastatic lesion that presented fainter, harder to segment cells. The ovarian sample can be seen to present both thin long cells as well as the standard circular cells seen in the BRCA2 samples.

**Figure 5.**
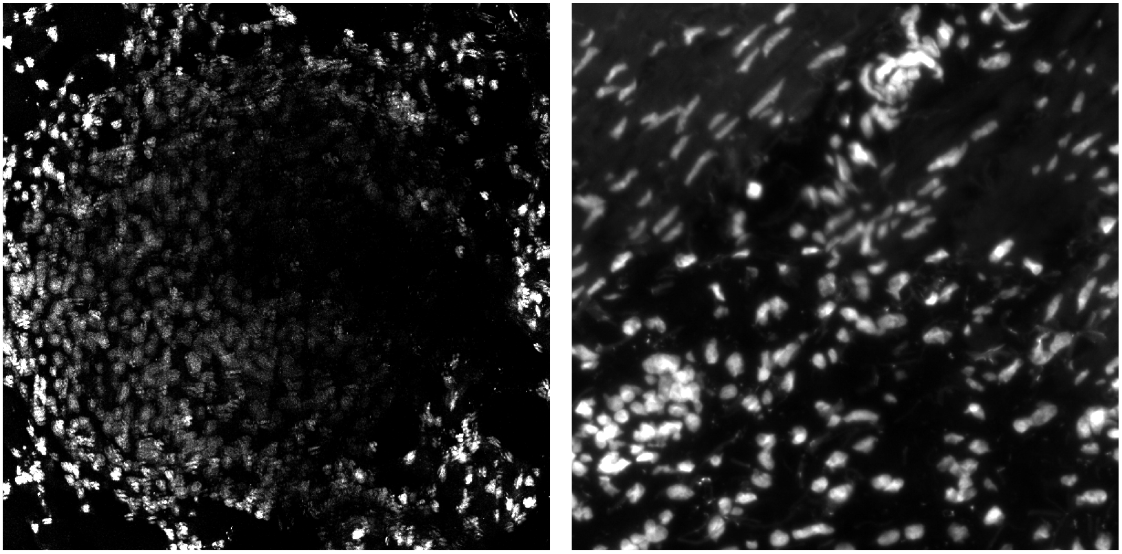
Two IMC images of mouse tissue; lung tissue on the left, ovarian tissue on the right.

In Figure 6 we focus on the metastatic lesion within the lung. For this image under-segmentation was a problem and so the uber mask was formed using the number of cell detections for the criteria of merit (neighborhood size of 40 pixels). The three algorithms that contributed to the majority of the uber mask detections were Cellpose v2, Mesmer and Watershed; the cell detections from these three methods as well as for the uber mask have been plotted as red contours over the input image. In Figure 7 we illustrate exactly how the uber mask is constructed by showing several cell detections within the lung sample. Within the region shown in Figure 7 the uber mask consists of cell segmentations from Cellpose v2, Mesmer and Watershed. The segmentations for Cellpose v2 are shown as purple contours overlaid onto the input image, the Mesmer segmentations are shown as teal contours and the Watershed segmentations are shown in light brown. The uber mask can be seen to be made up of contours from all of these methods; choosing Mesmer and Cellpose v2 over the Watershed technique for the specific areas where they detect a higher number of cells than the Watershed technique.

**Figure 6.**
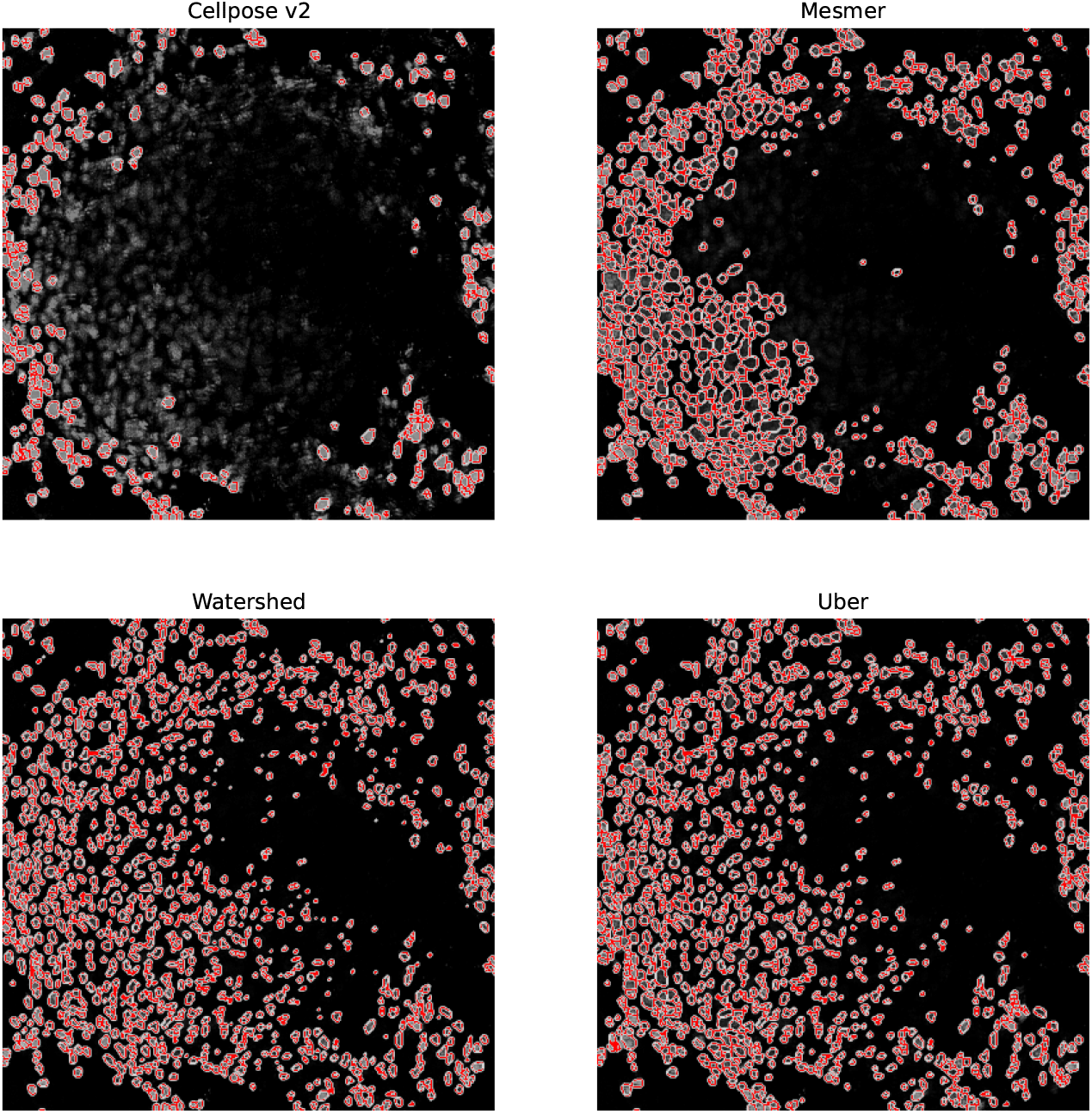
A specific region of the lung IMC sample. The cells segmented by Cellpose v2, Mesmer, Watershed and the uber mask are shown as red contours overlaid onto the nuclear channel data.

**Figure 7.**
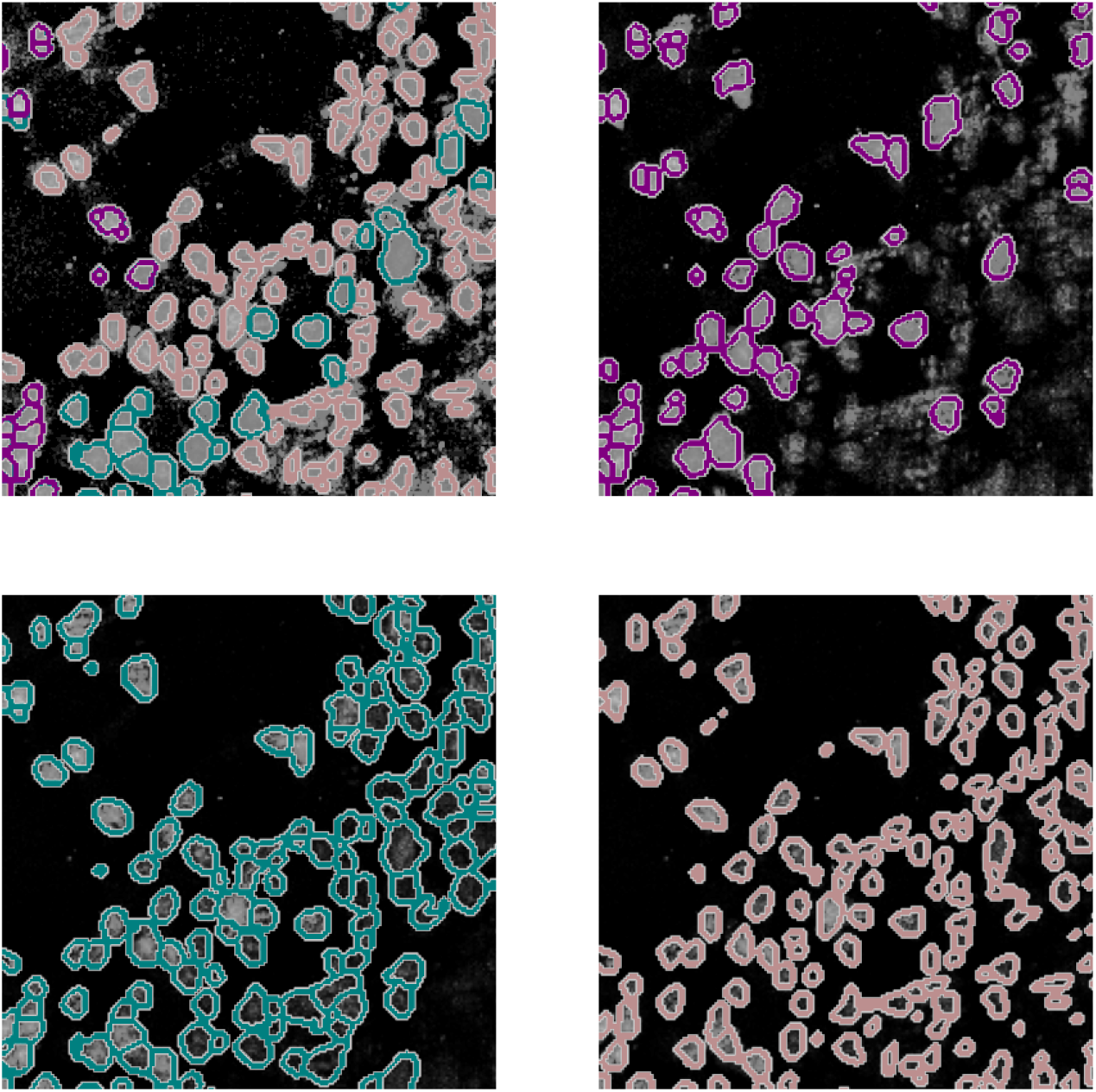
A specific region of the lung IMC sample. The top left image shows the uber mask cell detections as contours over the original image. The contours are colored according to the method that detected them. The top right, bottom left and bottom right images show the Cellpose v2, Mesmer and Watershed detections in purple, teal and light brown, respectively.

For Figure 8 we focus on a fairly typical region within the ovarian sample; this tissue sample was chosen as it contains both circular and elliptical shaped cells. For this sample under-segmentation was not seen to be problem, the difficulty for the segmentation algorithms was to be able to simultaneously identify two strikingly different cell structures. The uber mask was formed using the Jaccard score for the criteria of merit (neighborhood size of 40 pixels). The four algorithms that contributed to the majority of the uber mask detections were Cellpose, Cellpose v2, Mesmer and Mesmer v2; the cell detections from these four methods as well as for the uber mask have been plotted as red contours over the input image. For the specific region shown in Figure 9, the uber mask consists of cell segmentations from Cellpose, shown as light blue contours overlaid onto the input image, Cellpose v2 given as purple contours, Mesmer segmentations shown as teal contours and Mesmer v2 segmentations shown in light green. The uber mask can be seen to strike a balance between identifying all the cells clearly visible by eye, whilst not over segmenting and confusing background noise with genuine signal.

**Figure 8.**
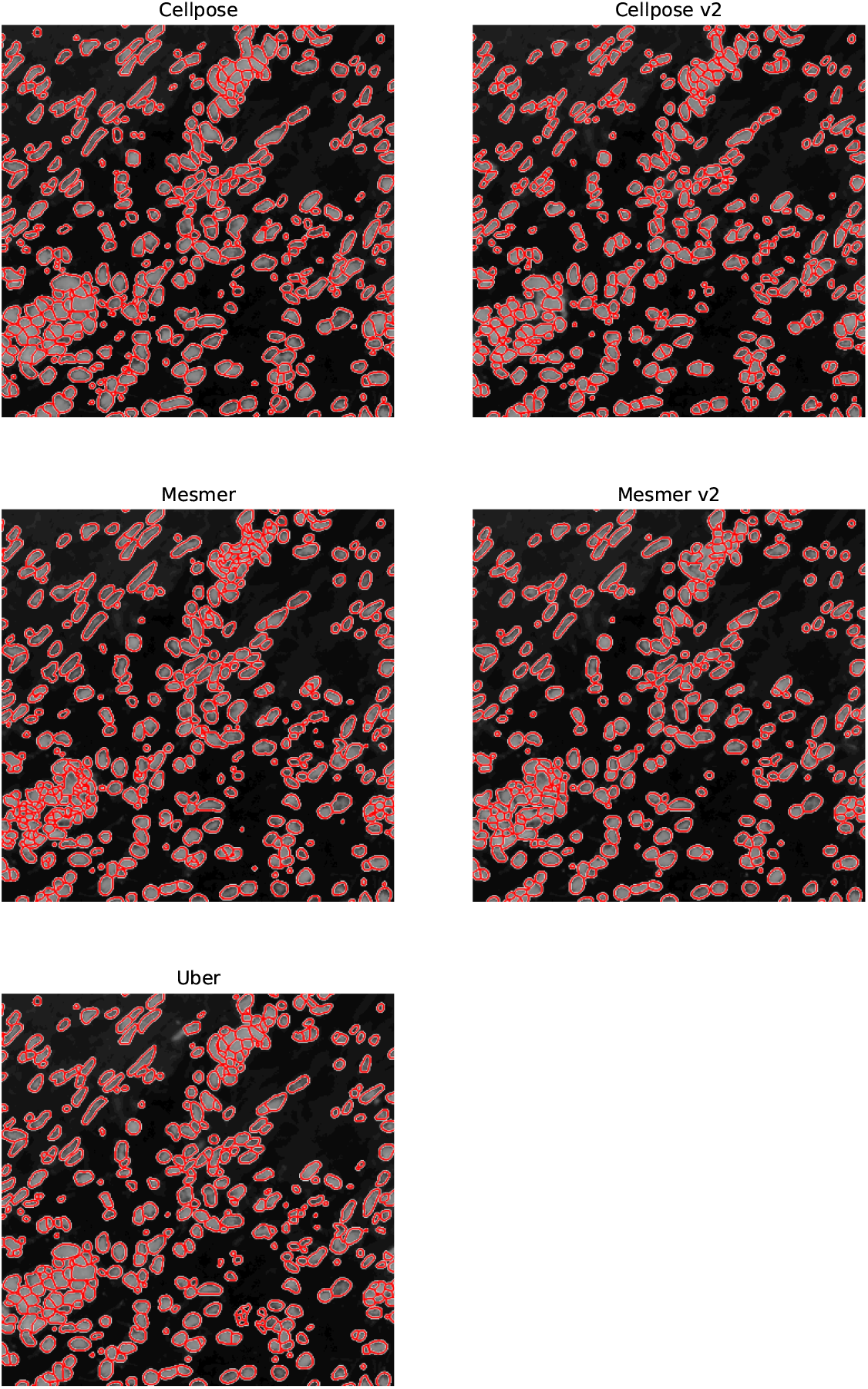
A specific region of the ovarian IMC sample. The cells segmented by Cellpose, Cellpose v2, Mesmer, Mesmer v2 and the uber mask are shown as red contours overlaid onto the nuclear channel data.

**Figure 9.**
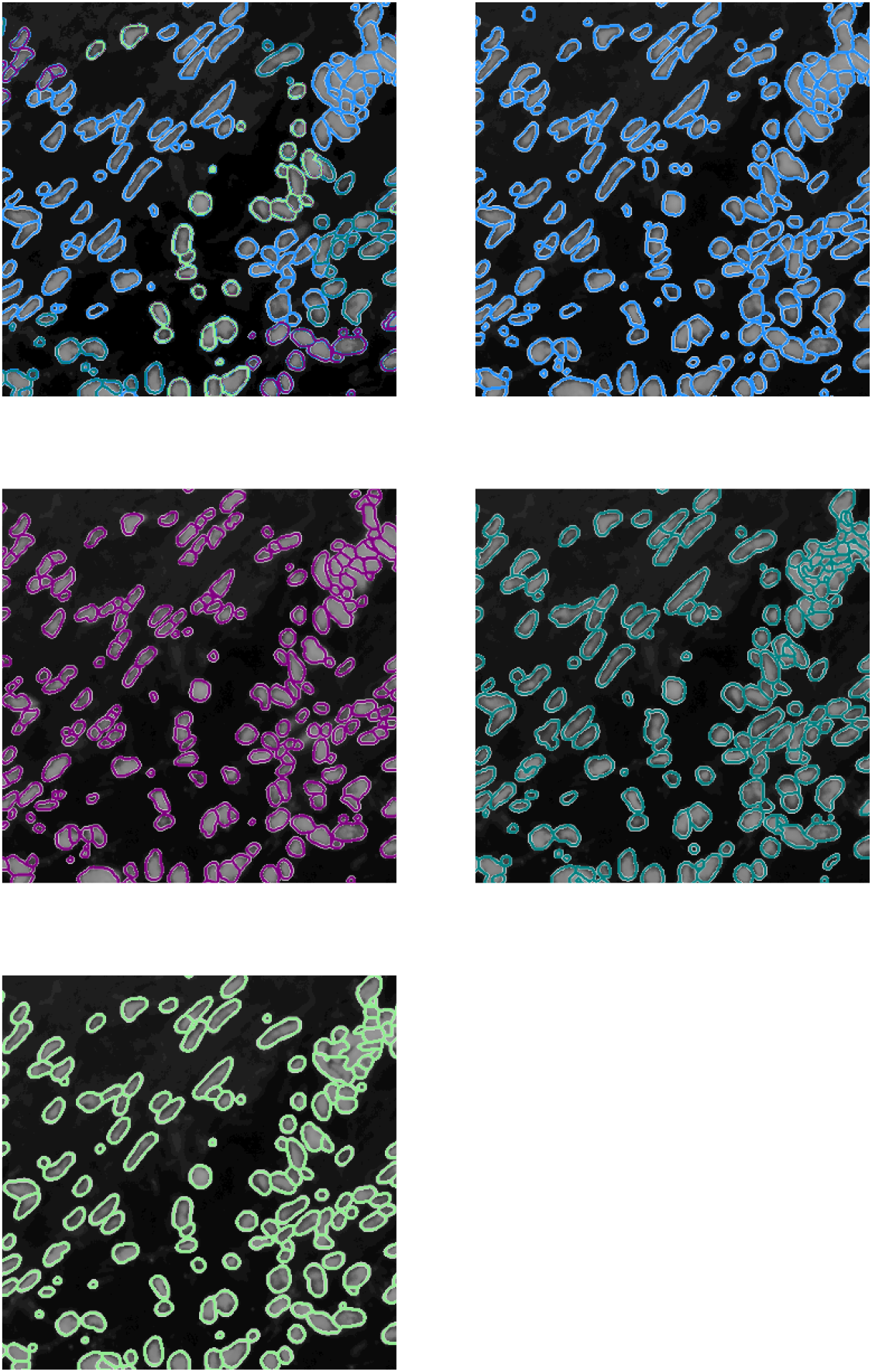
A specific region of the ovarian IMC sample. The top left image shows the uber mask cell detections as contours over the original image. The contours are colored according to the method that detected them. The top right, middle left, middle right and bottom left images show the Cellpose, Cellpose v2, Mesmer and Mesmer v2 detections in light blue, purple, teal and light green, respectively.

These two examples explicitly show the advantage of a consensus segmentation method with regards to optimally capturing as many genuine cell detections as possible whilst avoiding over-segmentation for a variety of cell types.

## 4 DISCUSSION

We have presented a new consensus voting technique which is capable of combing multiple segmentation algorithms for the segmentation of a single image. Our consensus voting is applicable to a wide range of spatial genomic imagery; we show results for IMC images in this study. Algorithms such as Cellpose and Stardist have been trained on both fluorescence and H&E images and so `CellSampler` can also be used for these types of images. The consensus technique presented in this work specifically divides the single image into local neighborhoods, allowing for different algorithms to lead the segmentation across different, localized regions. We have shown that for the BRCA2 dataset the consensus uber mask which votes based on the number of the cell detections within a local neighborhood produces the joint highest F1 and Jaccard scores, alongside the winning algorithms (Mesmer and Watershed for this analysis). We have also demonstrated the ability of the uber mask to capture the optimal number of cell detections across a selection of different IMC images presenting different cell types.

The `CellSampler` pipeline will optimize cell segmentation for large sample size image sets by enabling researchers to quickly test a variety of segmentation algorithms, identify the keys strength in what would appear to be the optimum method through a visual comparison of several segmented images and then construct a reliable segmentation for all of their hundreds/thousands of images through an uber mask optimized to make choices based on the user-selected key strength. This is marked improvement over the standard process of identifying one single preferred method through a quick visual inspection and then either relying on this single method, or painstakingly verifying by eye that it has performed optimally on each of the several hundred images.

As more segmentation techniques become publicly available they will be incorporated into `CellSampler` ensuring that the pipeline shown here continually evolves and maintains quality alongside the field’s latest cutting edge advances.

## AUTHOR CONTRIBUTIONS

M.O.I, E.A.G.S, T.W, A.M, M.A.S, N.A.W and D.B designed and built the `CellSampler` software. C.M.M and M.P.R performed the IMC experiments responsible for some of the data shown in this work as well, as running, testing and providing feedback necessary for the pipeline development of `CellSampler`. All authors read and approved the final manuscript.

## DATA AVAILABILITY STATEMENT

The `CellSampler` code will be made available at https://gitlab.developers.cam.ac.uk/astronomy/camcead/imaxt/public-code/cellsampler under a GNU General Public License version 3 (GPLv3).

## Footnotes

1 https://scikit-image.org/

